# IMPROVING MIN HASH VIA THE CONTAINMENT INDEX WITH APPLICATIONS TO METAGENOMIC ANALYSIS

**DOI:** 10.1101/184150

**Authors:** David Koslicki, Hooman Zabeti

## Abstract

Min hash is a probabilistic method for estimating the similarity of two sets in terms of their Jaccard index, defined as the ration of the size of their intersection to their union. We demonstrate that this method performs best when the sets under consideration are of similar size and the performance degrades considerably when the sets are of very different size. We introduce a new and efficient approach, called the *containment min hash* approach, that is more suitable for estimating the Jaccard index of sets of very different size. We accomplish this by leveraging another probabilistic method (in particular, Bloom filters) for fast membership queries. We derive bounds on the probability of estimate errors for the containment min hash approach and show it significantly improves upon the classical min hash approach. We also show significant improvements in terms of time and space complexity. As an application, we use this method to detect the presence/absence of organisms in a metagenomic data set, showing that it can detect the presence of very small, low abundance microorganisms.

## 1. Introduction

Min hash [3] is a fast, probabilistic method of estimating the Jaccard index, allowing for quick estimation of set similarity. Since the introduction of this technique by Broder (1997), this method has been used in a broad range of applications: from clustering in search engines [9], to association rule learning in data mining [8], to recent applications in computational biology [5, 19]. As a probabilistic method, min hash uses random sampling to estimate the Jaccard index of two sets. Bounds can be obtained on the probability of deviation from the true Jaccard index value in terms of the number of random samples used along with the magnitude of the true Jaccard value. Using the Chernoff bounds (Theorem 2.1), this probability of deviating from the true value grows exponentially as the size of the true Jaccard value decreases to zero. Hence, min hash returns tight estimates only when the true Jaccard index is large. This requirement for a large Jaccard index limits this technique to situations where the sets under consideration are of similar relative size and possess significant overlap with each other. In this manuscript, we introduce a modification of the min hash technique, which we call *containment min hash*, that does not possess this limitation and hence is appropriate for use when the sets under consideration have very different size.

After introducing the containment min hash technique, we derive rigorous probabilistic error bounds and compare these to those of the traditional min hash technique. This allows us to precisely state when the containment approach is superior to the classical approach. To demonstrate the practical utility of the containment min hash technique, we consider an application in the area of metagenomics (the study of all sampled DNA from a community of microorganisms), where the goal is to detect the presence or absence of a given genome in a metagenomic sample. This is an area of study where the sets of interest differ in relative size by orders of magnitude. This application highlights the improvements of this approach: in both theory and practice, in many situations of interest, containment min hash is significantly more accurate, has smaller computational complexity, and utilizes less memory than the traditional min hash approach.

We begin by giving a high-level summary of the main idea behind the containment min hash technique which is summarized in Figure 1. Consider the case of estimating the Jaccard index between two sets *A* and *B* of very different size. Briefly, the traditional min hash randomly samples from the union *A* ∪ *B* and uses the number of sampled points that fall in *A* ∩ *B* to estimate the Jaccard index. With more sampled elements falling in *A* ∩ *B*, the more accurate the Jaccard estimate will be. Part A) of Figure 1 demonstrates the case of sampling 100 random points from *A* ∪ *B* leading to 3 points lying in *A* ∩ *B*. In the containment min hash approach, we randomly sample elements only from the smaller set (in this case, *A*) and use another probabilistic technique (in this manuscript, a bloom filter) to quickly test if this element is in *B* (and hence in AnB). This is used to estimate the containment index, which is then used to estimate the Jaccard index itself. Part B) of Figure 1 demonstrates this approach while sampling only 50 points from *A* and finds 22 points lying in *A* ∩ *B*. This containment approach also experiences decreasing error as more points in *A* ∩ *B* are sampled, and so sampling from a smaller space (the set *A* instead of *A* ∪ *B*) leads to significant performance improvements.

**Figure 1.**
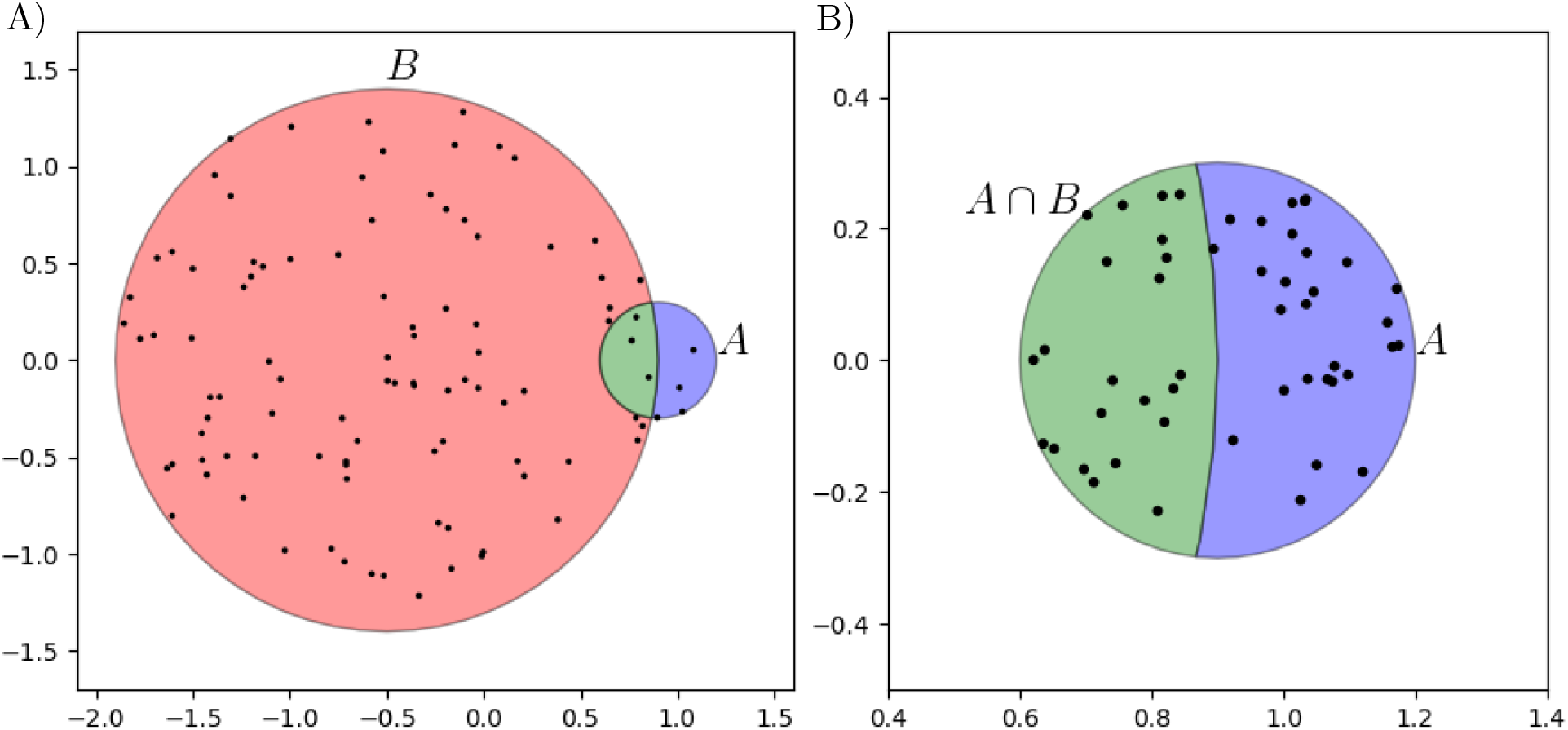
Conceptual comparison of classical min hash to the proposed containment approach when estimating the Jaccard index of very different sized sets. A) Sampling 100 points from *A* ∪ *B* (as is done in the classical min hash approach) leads to finding only 3 elements in *A* ∩ *B*. B) Sampling just 50 points of *A* and testing if a point *x* ∈ *A* ∩ *B*, finds 22 elements in *A* ∩ *B*. This latter approach will be seen to lead to a better estimate of the Jaccard index.

## 2. Methods

Before describing min hash and containment min hash, we recall a few definitions and results of interest.

### 2.1. Preliminaries

#### 2.1.1. Jaccard and Containment index

The Jaccard index, also known as the Jaccard similarity coefficient, measures the similarity of two sets by comparing the relative size of the intersection to the union [14]. That is, for two non-empty finite sets *A* and *B*,

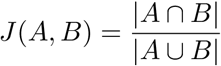

Hence, 0 ≤ *J (A, B)* ≤ 1 with larger values indicating more overlap.

To compare the relative size of the intersection to the size of *A*, we similarly define the containment index of *A* in *B* (both non-empty) as:

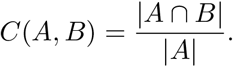

So 0 ≤ *C(A, B)* ≤ 1 and larger values indicate more of *A* lying in *B*.

#### 2.1.2. Chernoff Bounds

We will use the classic Chernoff bounds in its multiplicative form to estimate the probability of relative error in the probabilistic methods we consider.

##### Theorem 2.1

([18, Thm 4.4-4.5]). *Suppose X_1_, X_2_, …,X_n_ are independent, identically distributed Bernoulli random variables and let* 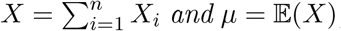, *then the following statements hold for* 0 < δ < 1:

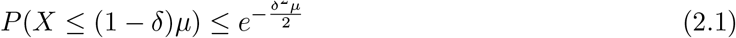

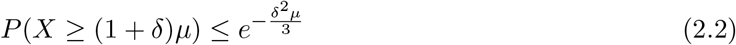

*And hence*,

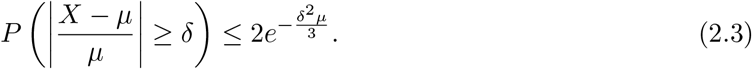

#### 2.1.3. Bloom Filters

As mentioned in the Introduction, we will need a fast test of set membership to implement the containment min hash approach. While there exist many different such data structures (such as Bloom Filters [2], skip lists [21], cuckoo filters [10], quotient filters [1], etc.), we utilize the Bloom filter due to its ubiquity [6,7,12,17,19,20,22,23] in the application area considered here (computational biology) and the ease of its mathematical analysis.

We give a brief description of the construction of a bloom filter following the exposition of [18]. Given a set *B* with cardinality *n* = |*B*|, a bloom filter 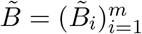 is a bit array of length *m* where *m* is a chosen natural number. Fix a set of hash functions *h_1_,…,h_k_* each with domain containing *B* and with range {1, …, *m*}. Initially, each entry in the bloom filter is set to zero: 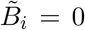 for each *i* = 1, …, *m*. For each *x* ∈ *B* and *j* = 1, …, *k*, we set 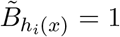. Given an element *y* in the domain of our hash functions, if

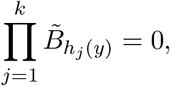

then by construction, we know that *y* ∉ *B*. Because of this, we write 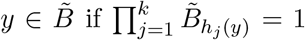, and 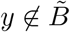 otherwise. A straightforward calculation (with idealized hash functions) shows that the probability of 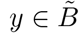 and yet *y* ∉ *B* is given by:

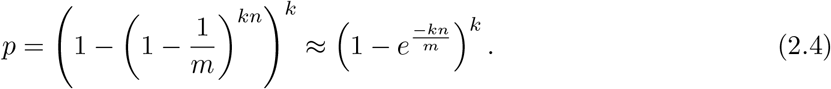

The optimal false positive rate *p* is given by 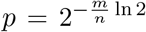 when the number of hash functions is given by 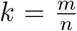. Conversely, given a target false positive rate *p*, the optimal length *m* of the bloom filter is given by 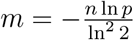.

### 2.2. Containment Min Hash

Before detailing the containment min hash approach, we recall the classic min hash for comparison purposes.

#### 2.2.1. Classic Min Hash

By the *classic min hash*, we mean the construction of Broder (1997). We detail this construction now. Given two non-empty sets *A* and *B*, we wish to estimate *J(A,B)*. Fix 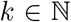 and select a family of (min-wise independent [4]) hash functions {*h*_1_,…, *h_k_*} of 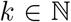 each with domain containing *A* ∪ *B*. For a set *S* in the domain of the hash functions, define 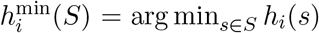 as an element of *S* that causes *h_i_* to achieve its minimum value on *S*. Given appropriate hash functions (or an order on *S*), this minimum is unique. Define the random variables

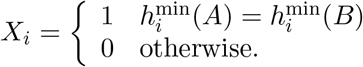

The probability of a collision (that is 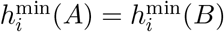 and hence *X_i_* = 1) is equal to the Jaccard index of *A* and *B* [4] and hence the expectation is given by

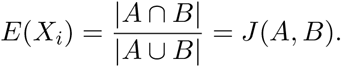

Thus, for 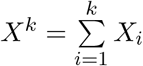, the expectation is given by *E(X^k^) = kJ(A, B)* and so 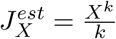 is used as the estimate of *J (A, B)*. Note that in practice, a single hash function *h* is commonly used and the elements hashing to the smallest k values are used in place of the *h_i_*.

Applying the two-sided Chernoff bounds from equation (2.3) to the classic min hash approach, we have

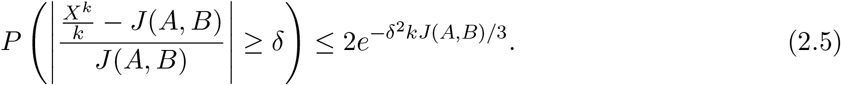

Thus, two quantities control the accuracy of this method for δ fixed: *k* and *J (A,B)*. Setting *t* as the threshold of probability of deviation from the Chernoff bounds (i.e. *t* = 2*e*^-δ^2^*kJ*(*A,B*/3^), we calculate the number of hash functions *k = k_X_* required to achieve this threshold:

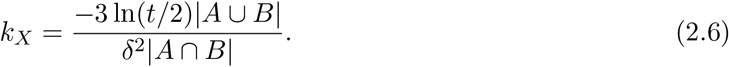

#### 2.2.2. Containment Min Hash

The containment min hash approach we propose differs from the classic min hash in that the family of *k* hash functions {*h*_1_,…, *h_k_*} have domain containing *A* and we randomly sample from *A* instead of *A* ∪ *B*. This results in estimating the containment index *C = C (A,B)* instead of the Jaccard index *J = J (A,B)*, but we will late show how to recover an estimate of *J (A, B)*. The containment approach proceeds as follows: Let 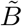 be a bloom filter with given false positive rate *p* and define the random variables

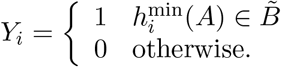

These random variables essentially sample (uniformly) randomly from *A* and tests for membership in w*B* via the bloom filter 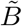. Following the same proof of Broder (1997), it is straightforward to show that 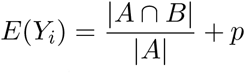. Thus for 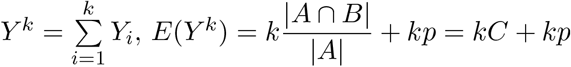. Hence, 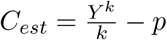 as the estimate of *C(A, B)*.

Applying the two-sided Chernoff bounds from equation (2.3) now gives

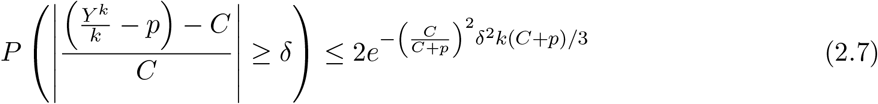

It is important to note that 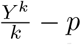 estimates *C(A, B)* and not *J(A, B)*, as desired. To directly compare these quantities, we must derive the Jaccard index from the containment estimate and calculate the probability of deviation from the true value of the Jaccard index. To that end, for *C_est_* the estimation of the containment, let 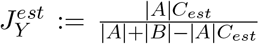. Note that in practice, a fast cardinality estimation technique (such as HyperLogLog [11]) can be used to approximate |*A*| and |*B*|. The bound in the following proposition can be directly compared to that of equation (2.5).

##### Proposition 2.2.

*For* 0 < δ < 1, *let* 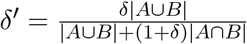, *then*

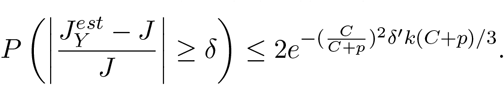

##### Proof.

We first derive a useful characterization of the probability under consideration:

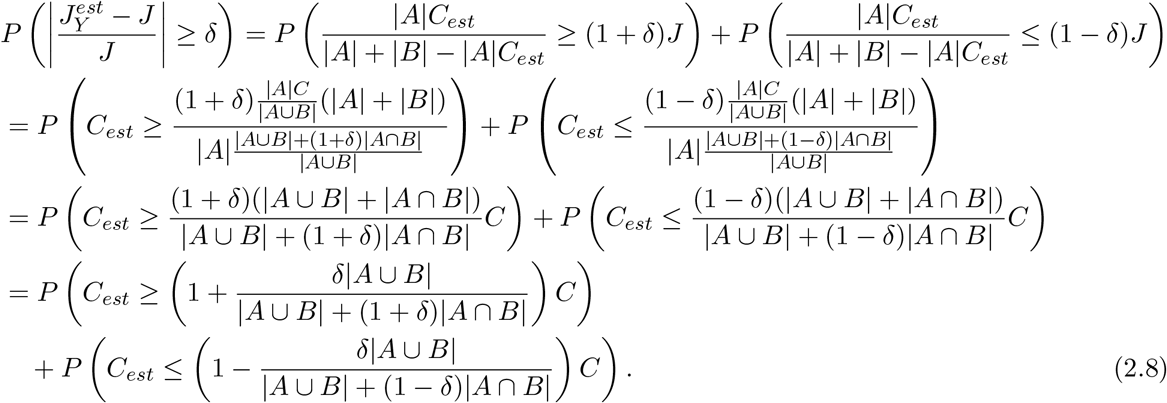

Then for δ′ as defined in the statement of the proposition and for 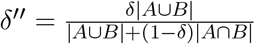, note that 0 < δ′,δ′″ < 1. Then using the two-sided Chernoff bounds from equations (2.1) and (2.2) applied to equation (2.8), we have that

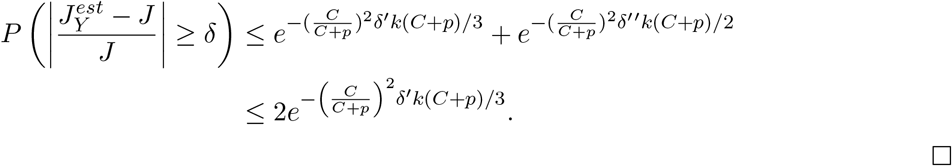

□

Figure 2 gives a comparison of the bounds of deviation (as a function of δ for a fixed number of hash functions) for the classical min hash Jaccard estimate (equation (2.5)) and the containment min hash estimate of the Jaccard index (Proposition 2.2). This figure shows that the containment min hash approach has a significantly smaller probability of the estimate deviating from the true value.

**Figure 2.**
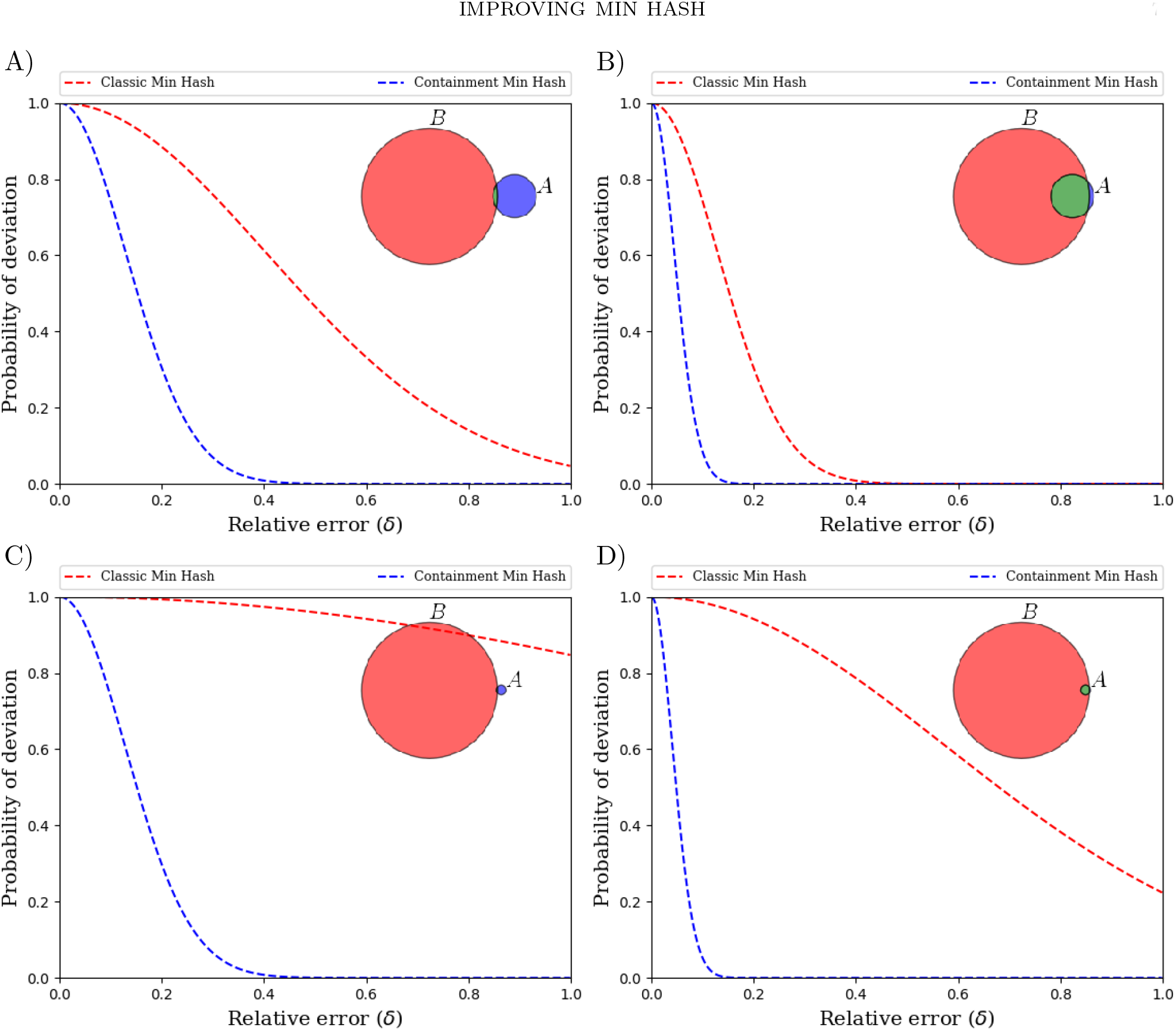
Comparison of the probability of deviation (red line: equation (2.5), blue line: Proposition (2.2)) versus the relative error δ with a fixed number of hash functions (1,000). Relative sizes and Jaccard indexes of the sets in A)-D) are overlain as colored discs. Throughout, the containment min hash estimate of the Jaccard index has a much lower probability of error than the classical min hash approach. Note that in part D), there is approximately an 80% chance to have a deviation greater than 0.4 in the classic approach, whereas in the containment approach, the chance of having a deviation of greater than just 0.2 is almost zero.

Now, let *k = k_γ_* be the number of hash functions which is required to achieve a desire threshold upper bound of 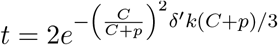. We then have that

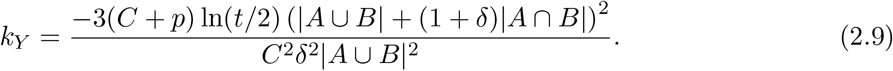

## 3. Results

We begin by detailing the theoretical results obtained before turning to the application of interest.

### 3.1. Theoretical Results

#### 3.1.1. Number of Hash Functions Required

For both the classical min hash approach and the containment min hash approach, we use equations (2.6) and (2.9) to compare the number of hash functions *k_X_* and *k_Y_* required for a specified threshold of probability of deviation *t*. Calculating, we obtain:

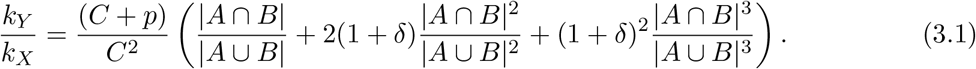

Note that 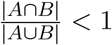, so we have that:

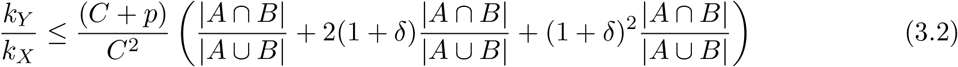

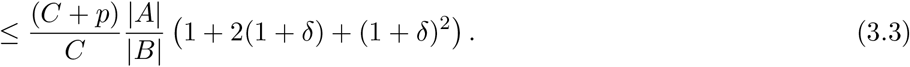

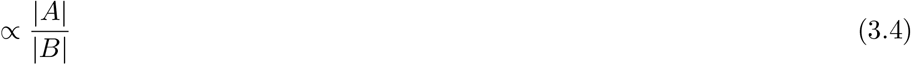

where the last proportionality holds when *C* is bounded away from zero. Hence, the containment approach uses a fraction (proportional to 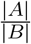) of the hashes that the classical approach uses to achieve the same bound of error. When |*B*| is significantly larger than |*A*|, this reduction in the number of hash functions required can be significant. In Figure 3, we compare the relative error δ in estimating the Jaccard index to the number of hash functions required for the classical and proposed approach to have a probability of ≤ 1% of more than δ relative deviation in their estimate. Interestingly, the number of hashes required by the containment approach is nearly constant as a function of 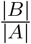 whereas for the classical approach, the number of hashes required is increasing. Figure 4 depicts this observation.

**Figure 3.**
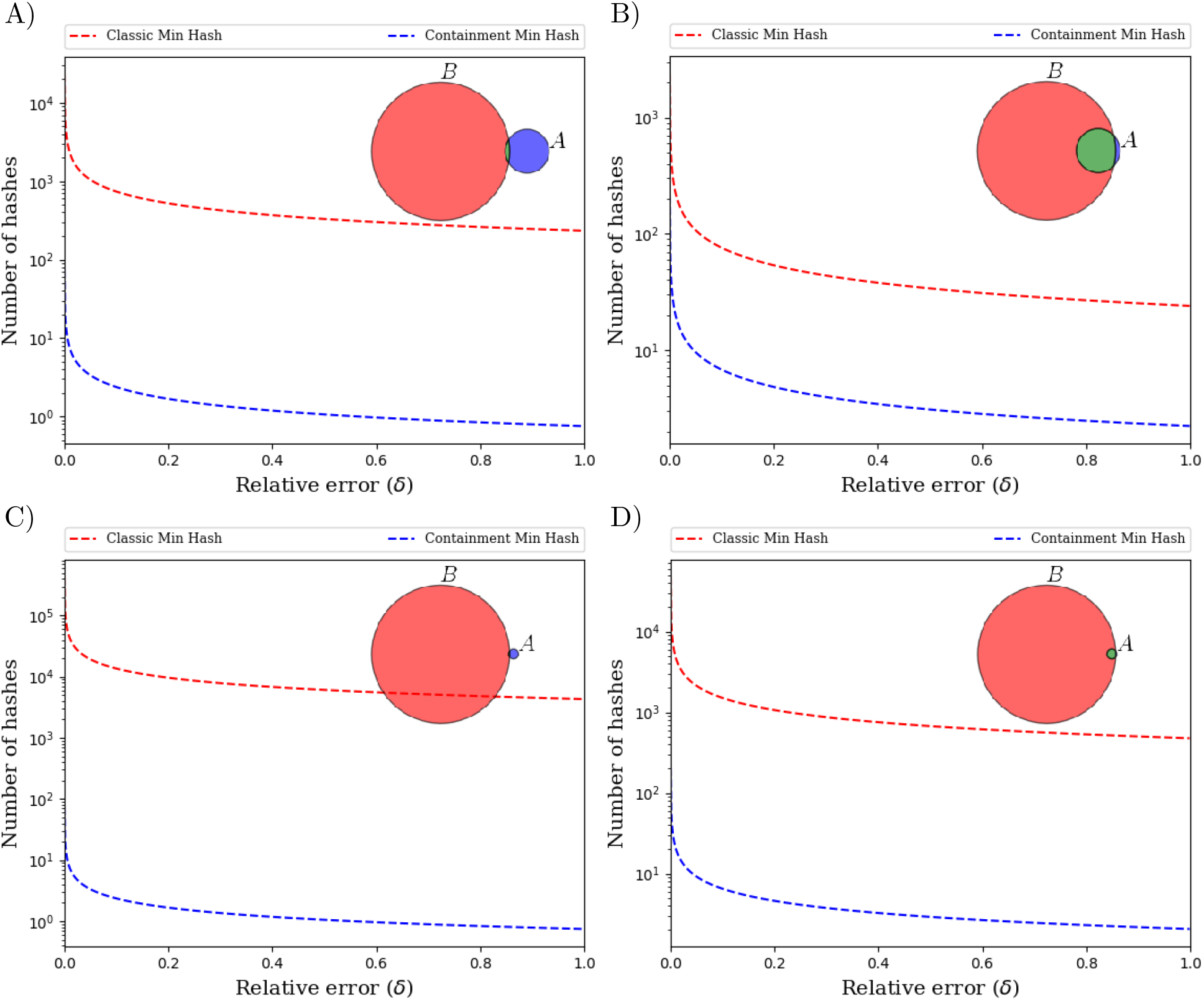
Comparison of the number of required hashes and relative error δ with probability of deviation ≤ 1% := *t*. The relative sizes and Jaccard indexes of the sets in A)-D) are overlain pictorially as colored discs. In all cases, the containment min hash estimate of the Jaccard index uses significantly fewer hash functions. For example, in part D), the classic min hash method (red line) needs ≈ 35, 339 hash functions to have less than 1% chance to have greater than 1% relative deviation in its estimate, whereas with the same thresholds, the containment min hash estimate of the Jaccard index (blue line) needs only ≈ 152 hash functions.

**Figure 4.**
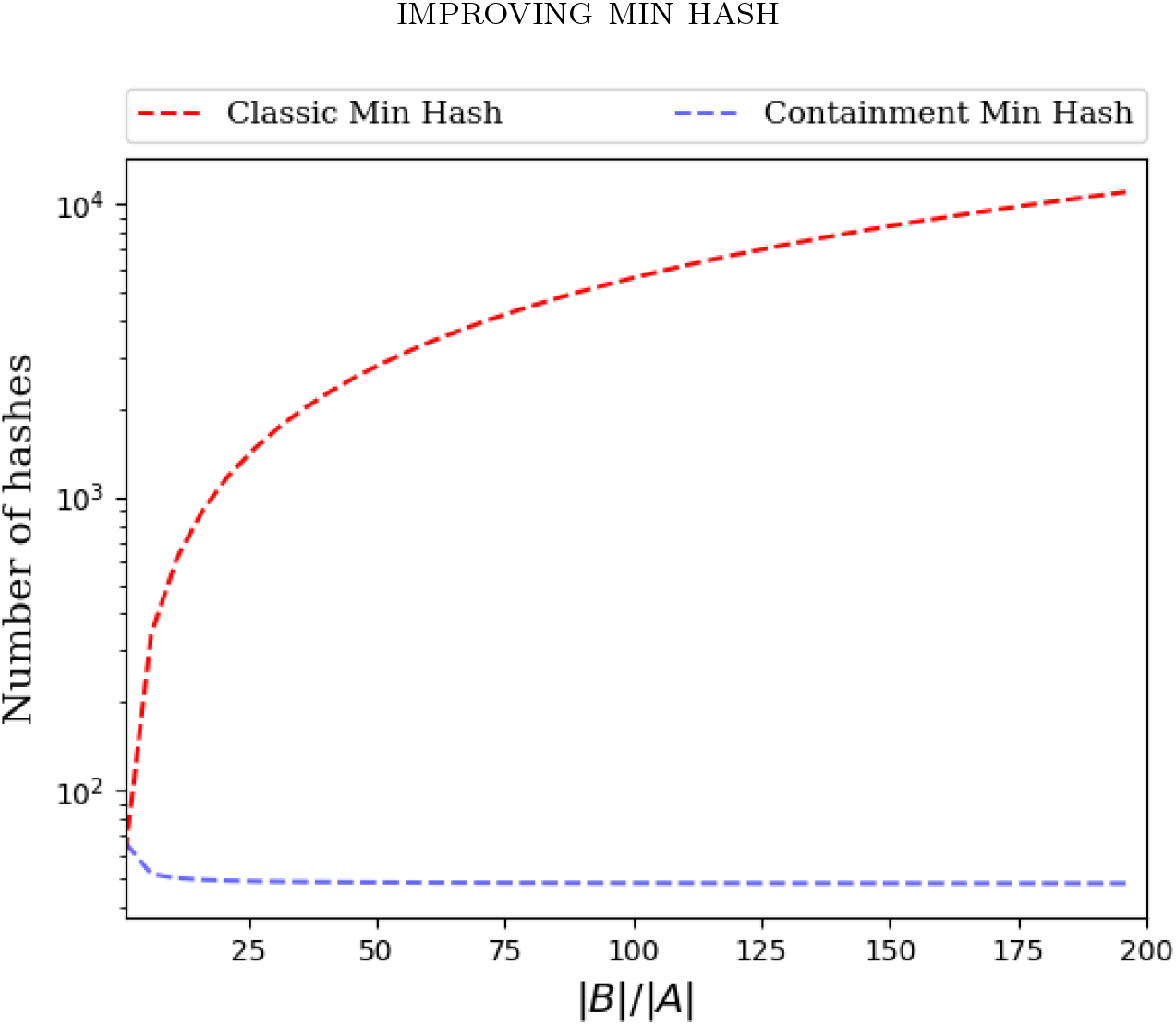
Comparison of the number of hash functions required by each method as a function of the relative sizes of the sets under consideration. The relative error δ = 0.1 and the probability of deviation *t* = 0.01 are fixed. Increasing 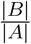, the number of required hash functions required for the classic min hash approach is increasing (red line). However, for the containment approach, the number of hash functions is nearly constant for 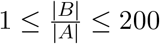 (blue line) which is in agreement with equation 3.4.

#### 3.1.2. Time/Space Complexity

Given sets *A* and *B* of size *m* and *n* respectively, both the classic min hash and containment min hash require 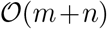 time to form their respective data structures. When calculating their estimates of the Jaccard index *J (A, B)*, both approaches use computational time linear in the number of hash functions required (in the case of the containment min hash approach, this is due to Bloom filter queries being constant time [2]). Because of equation (3.1) and the discussion that followed, this implies that the ratio of time complexity of the classical min hash to the containment min hash is 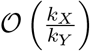. Given equations (3.1) and (3.4), when *B* is very large in comparison to *A*, this implies that the containment min hash approach is significantly faster than the classical min hash approach.

In terms of space complexity, if all one desires is an estimate of the Jaccard index from a single pair of sets, the containment approach will use more space as a bloom filter must be formed from one of the sets. However, in the application of interest, we have one large (*n* ≫ m) “reference” set *B* and many smaller sets 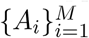 each with |*A_i_*| ≤ *m*. Here, we wish to estimate *J(B, A_i_)* for each *i*. In this case, the additional space required to store a bloom filter is dwarfed by the space savings that come from using significantly fewer hash functions. Indeed, for *S_X_* and *S_Y_* the space required for the classic and containment min hash respectively, we have that *S_X_ ∝ k_X_*(*M*, 1) and *S_y_* ∝ *k_Y_* + 1.44 log_2_(1/*p*)*n* where the proportionality is in terms of the number of bits required to store a single hash value. Holding this constant fixed (as well as *p* and *t*), we then have:

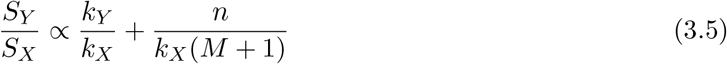

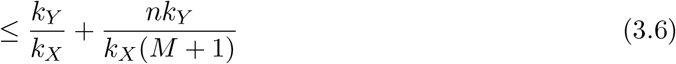

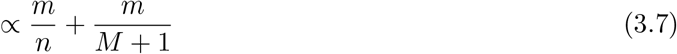

demonstrating that for large enough *M*, the containment approach will use less space than the classical min hash approach. A more detailed space analysis can be obtained by combining equations (2.6) and (2.9). For example when |*B*| = 10^8^ and one wishes to estimate the Jaccard index of *M* = 30000 smaller sets *A_i_* with containment *C(A_i_, B)* ≥ 80% that are small enough that *J(A_i_, B)* ≤ 10^-4^ and with *t* = 95% confidence that the estimate is not off by more than δ = 50%, using a false positive rate of *p* = 0·01 for the bloom filter results in a 33X space savings when using the containment approach instead of the classical min hash approach.

### 3.2. Simulated and Real Data

In this section, we compare the classic min hash approach to the proposed containment method on real and simulated data. All code and software required to reproduce the results contained here are available at:

~~~
https://github.com/dkoslicki/MinHashMetagenomics.
~~~

Included in this repository is an automated script that can reproduce this paper in its entirety:

~~~
https://github.com/dkoslicki/MinHashMetagenomics/blob/master/src/MakePaper.sh
~~~

#### 3.2.1. Simulated data

Here we illustrate the improved accuracy of the containment min hash approach over classical min hash in estimating the Jaccard index. To that end, we generated two random strings *w_A_* and *w_B_* on the alphabet {*A, C, T, G*}. We set |*w_A_*| = 10,000 and |*w_B_*| = 15 to simulate the situation of interest where one wishes to estimate the Jaccard index of two sets of very different size. We picked a *k*-mer (substring of length *k*) size of 11 and considered the sets *A* and *B* to be the set of all *k*-mers in *w_A_* and *w_B_* respectively. We then generated strings *w_C_i__* of increasing length, formed a set *C_i_* of all its *k*-mers, and considered *J* (*A* ∪ *C_i_, B* ∪ *C_i_*). The number of hash functions for each method was fixed to be *k_x_ = k_Y_* = 100. Figure 5 depicts the comparison of the containment min hash approach with the classical min hash Jaccard estimate on this data and effectively illustrates the results in section 2.2 showing improved performance of the containment min hash approach. The mean and variance of the classic min hash approach on this data was –0.001461 ± 0.001709 while using the containment approach was 0.000818 ± 0.000007, demonstrating a substantial decrease in variance. This improved variance was observed over a range of *k*-mer sizes, number of hashes, and lengths of input strings.

**Figure 5.**
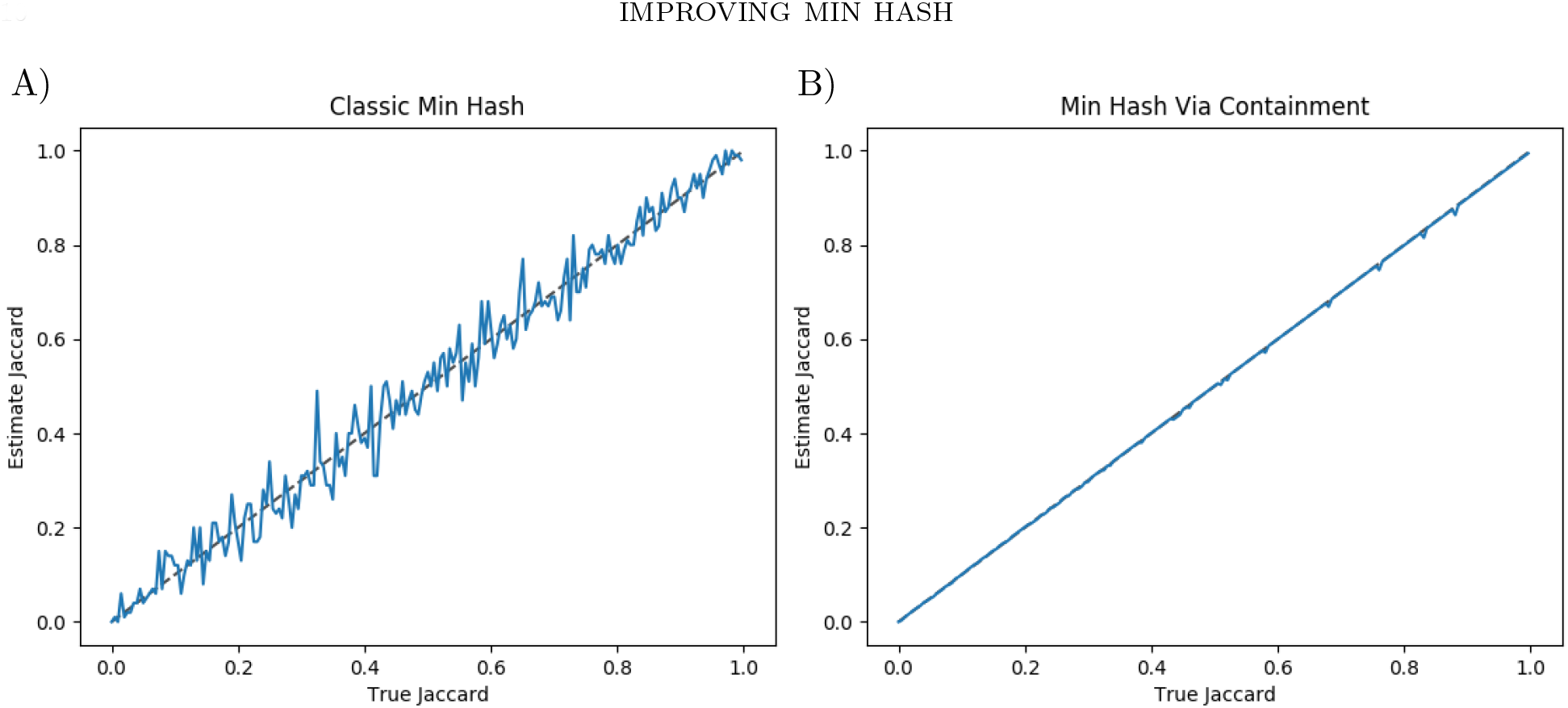
Comparison of the containment min hash approach to the classical min hash estimate of the Jaccard index on synthetic data. Each method utilized the 100 smallest hashes of the murmer3 hash function on the 11-mers of two randomly generated strings with sizes 10,000 and 15 respectively after appending a common substring of increasing size. A) Classical min hash estimate of the Jaccard index. B) Containment min hash estimate of the Jaccard index.

#### 3.2.2. Simulated biological data

To demonstrate the exponential improvement of the containment min hash approach over the classical min hash for increasing sample sizes, we contrast here the mean relative performance of each approach on simulated biological data. We utilized GemSIM [16] to simulate two sets of metagenomic data from randomly selected bacterial genomes. We them aim to estimate the Jaccard index between the metagenomic data and each of the bacterial genomes (thereby simulating the case when attempting to detect if a bacterial genome appears in a given metagenomic sample).

For the first set of simulated data, we used GemSIM to simulate 10K reads (of length 100) from 20 randomly selected bacterial genomes *G_i_* (considered here as the set of all *k*-mers in the genome). We fixed the *k*-mer size to *k* = 11. We then formed a set *M*_1_ of all 11-mers in all reads in the simulated metagenome. We then repeated this 16 times. A false positive rate of 0·001 was used for the bloom filter in the containment approach.

The second set of simulated data was produced similarly, except for the fact that we used 1M reads of the same length as before and formed a set *M*_2_ of all 11-mers in all reads of the metagenome. Again, we then repeated this 16 times using the same false positive rate for the bloom filter.

Figure 6 depicts the relative error of the classic min hash approach and the containment approach on these two sets of simulated data when estimating *J* (*M*_1_, *G_i_*) in part A), and *J* (*M*_2_, *G_i_*) in part B), as a function of the number of hashes used. Observe that the containment approach has significantly less error when, as is commonly seen in practice, the number of 11-mers in the sample is appreciable in comparison to the number of 11-mers in a given reference genome *G_i_*. This improvement of the containment approach over the classic approach continues to grow as the metagenome size grows in relation to the reference genome sizes.

**Figure 6.**
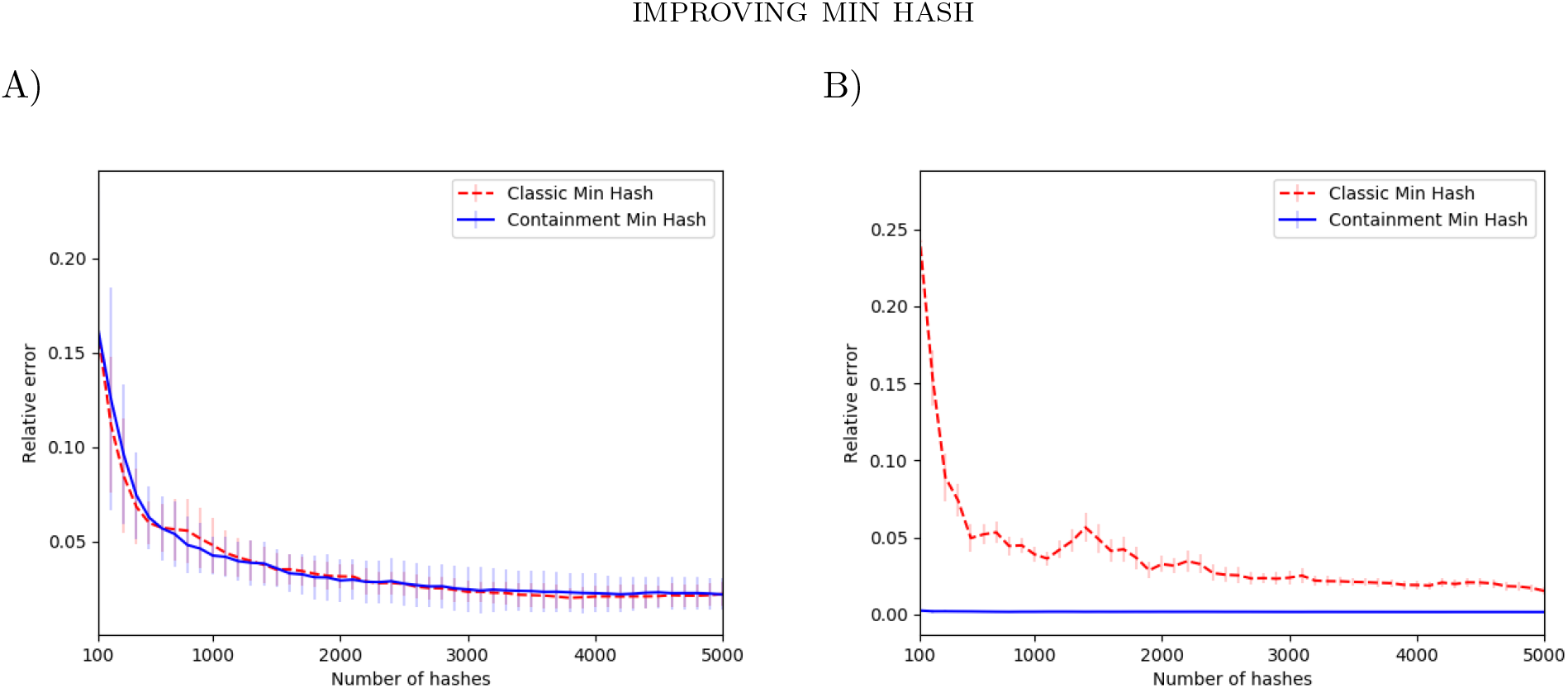
Comparison of the relative error of the containment min hash approach to the classical min hash estimate of the Jaccard index on simulated biological data. a) On 16 replicates of samples consisting of 20 genomes *G_i_* with only 10K reads, showing the similarity of the methods in estimating *J*(*M*_1_, *G_i_*) when the sets to be compared are roughly the same size. b) On 16 replicates of samples consisting of 20 genomes *G_i_* with 1M reads, demonstrating the improvement of the containment approach in estimating *J*(*M*_2_, *G_i_*) when the sets are of significantly different size.

### 3.3. Real biological data

Real metagenomes contain many magnitudes more *k*-mers than those found in any reference organisms, indicating the advantage of the containment min hash approach to determining the presence/absence of reference organisms in a given metagenome. To evaluate this, we analyzed a subset of DNA generated by the study in [13] consisting of those reads contained in the sample 4539585.3.fastq. This sample consisted of 25.4M reads with average length of 65bp. We formed a bloom filter consisting of all 21-mers of this sample and formed 500 hashes from each of 4,798 viral genomes obtained from NCBI [24]. Utilizing the proposed containment min hash approach, we found the largest containment index between the reference viral metagenomes and the sample to be 0.0257 for the virus *Sauropus leaf curl disease associated DNA beta* which corresponds to a Jaccard index of 2.398e-08. Note that with this small of a Jaccard index, the results in Section 3.1.1 show that the classic approach would need millions of hash functions to accurately estimate this quantity.

To evaluate if this extremely low-abundance organism is actually present in the sample, we utilized the SNAP alignment tool [25] to align the sample to the *Sauropus leaf curl disease associated DNA beta* genome. The script *MakeCoveragePlot.sh* provides the exact commands and parameters used to perform the alignment. We found that 288 reads aligned with a MAPQ score above 20 (i.e. high-quality alignments). The coverage of the viral genome is depicted in Figure 7 using a square-root scale and a window size of 10. These high-quality mapped reads to such a small genome lends evidence to support the claim that this particular virus is actually present in the sample metagenome.

**Figure 7.**
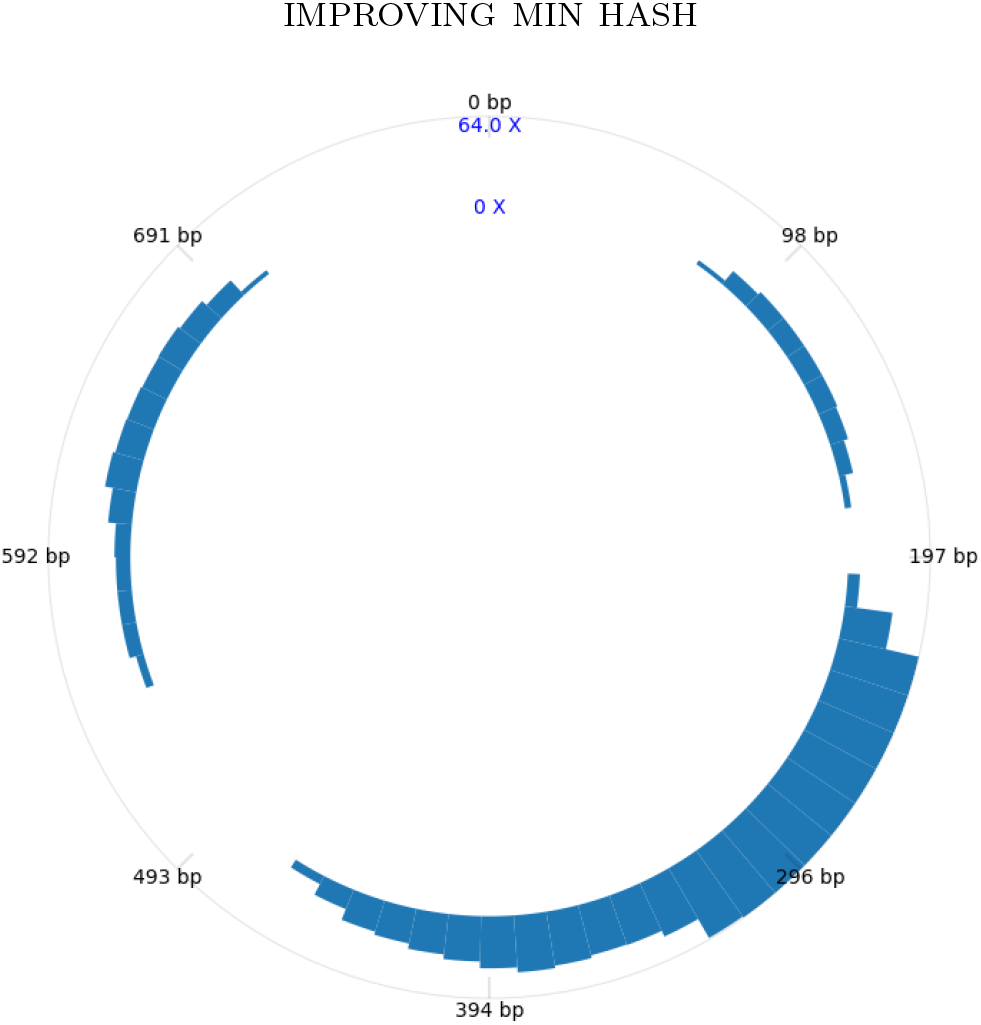
Plot of the real metagenomic sample alignment coverage to the virus *Sauropus leaf curl disease associated DNA beta* detected by the proposed containment min hash approach. A total of 288 reads aligned with a MAPQ score above 20 (i.e. high-quality alignments) using the SNAP aligner [25]. A square root scale and a window size of 10 was used for the plot, resulting in an average per-window coverage of 24.217X.

## 4. Conclusion

In this manuscript, we introduced “containment min hash”: an improvement to the min hash approach of estimating the Jaccard index. We derived exact formulas for the probability of error for this method and also showed that it is faster, more accurate and uses less space in many situations of practical interest. This improved approach was used to analyze simulated and real metagenomic data and was found to give results superior to those of the classic min hash approach. As the theoretical results we obtained are agnostic to the application of interest, we believe this method will be useful in any application where estimates of similarity of very differently sized sets is desired. Hence, this advancement can be useful outside of the field of metagenomics and computational biology, with possible applications in data mining, web clustering or even near duplicate image detection.

